## Supplemental doc for "Two dramatically distinct archaeal type IV pili structures formed by the same pilin"

### Supplementary Material

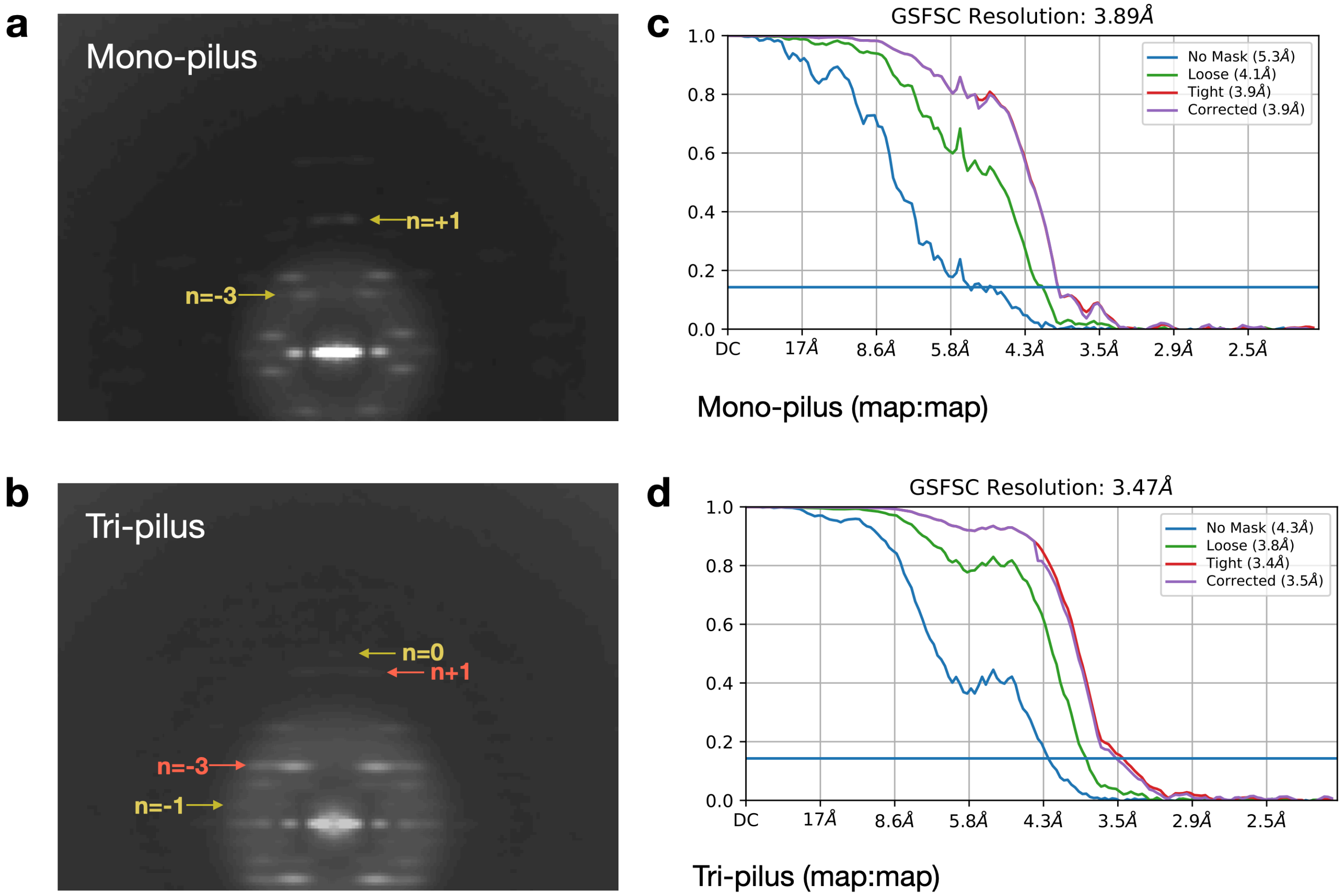


**Supplementary Figure 1. Average power spectrum and Fourier Shell Correlation (FSC) calculations of mono-pilus and tri-pilus**

Average power spectra of particles from 2D class average of mono-pilus (**a**) and tri-pilus (**b**). The Bessel orders of the layer lines used to determine the global helical symmetry are labeled in yellow. For tri-pilus, the layer-lines corresponding to the features generated from the inner helices are labeled in red. The map:map "gold-standard FSC" curves are shown for the mono-pilus (**c**) and tri-pilus (**d**).

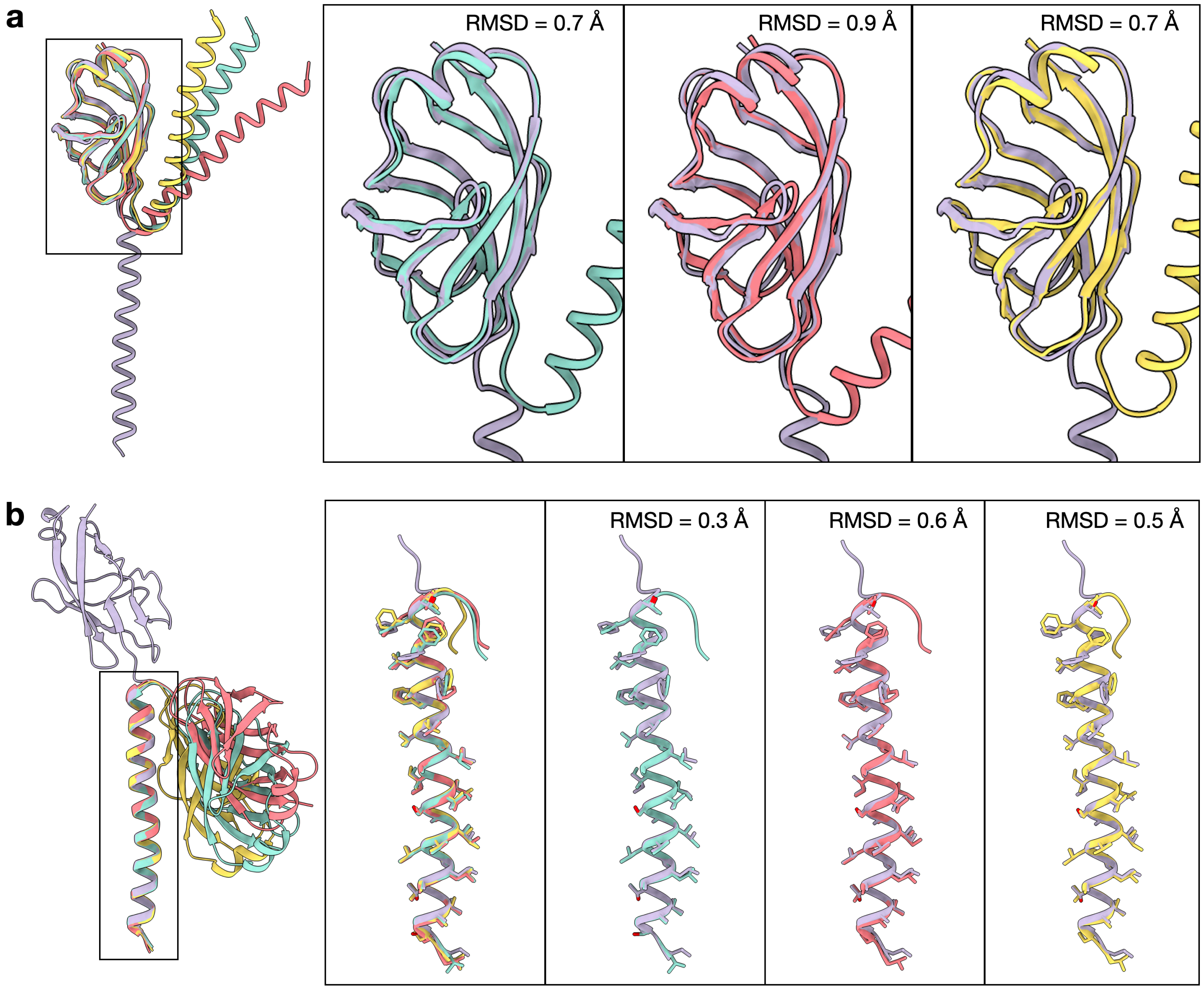


**Supplementary Figure 2. Alignment of C-terminal and N-terminal domains of mono-pilin and tri-pilins**

(**a**) Alignment of the C-terminal Ig-like globular domains is depicted, comparing the mono-pilin (lavender) with each of the three tri-pilins (A, cyan; B, red; C, yellow). Insets provide a magnified view of region (**a**), focusing on the mono-pilin and a single tri-pilin subunit, along with the labeled backbone RMSD (Root Mean Square Deviation).

(**b**) Alignment of each of the three tri-pilins to the mono-pilin by the N-terminal long helix, following the same visualization approach as in panel (**a**). Sidechains for residues 13-45 are additionally shown.

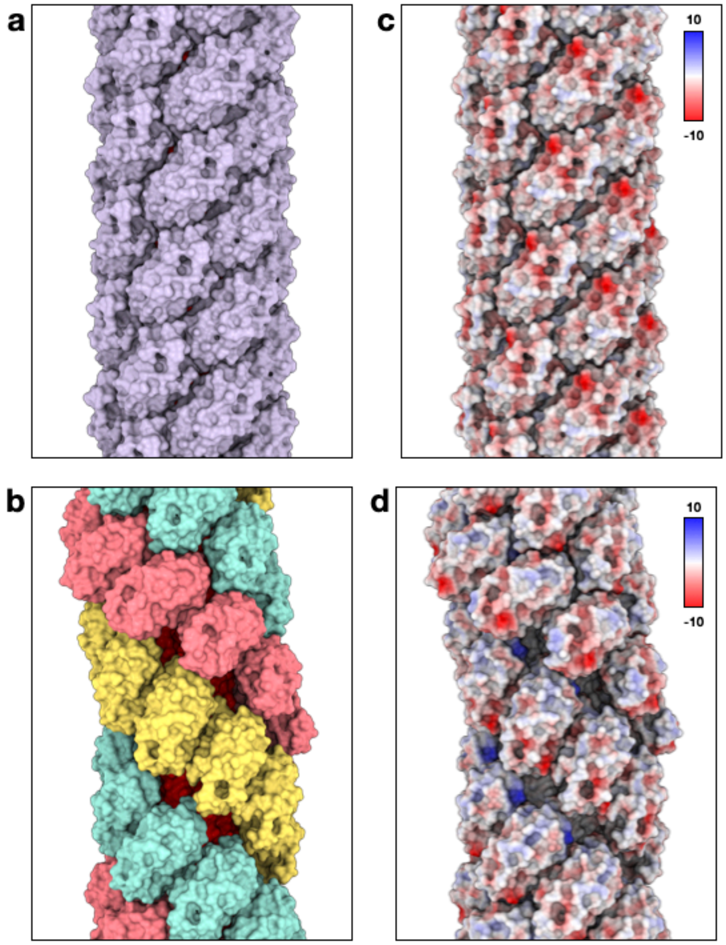


**Supplemental Figure 3. Solvent-excluded and electrostatic surfaces of the mono-pilus and tri-pilus**

(**a**) Molecular solvent-excluded surface of the mono-pilus. The inner helices are colored in red.

(**b**) Molecular solvent-excluded surface of the tri-pilus. The inner helices of all pilins are colored in red.

Electrostatic surface of the mono-pilus (**c**) and the tri-pilus (**d**). Scale bar displays the color-coding of the electrostatic potential in units kcal·mol$^{-1}$·$e^{-1}$ at 298 K.

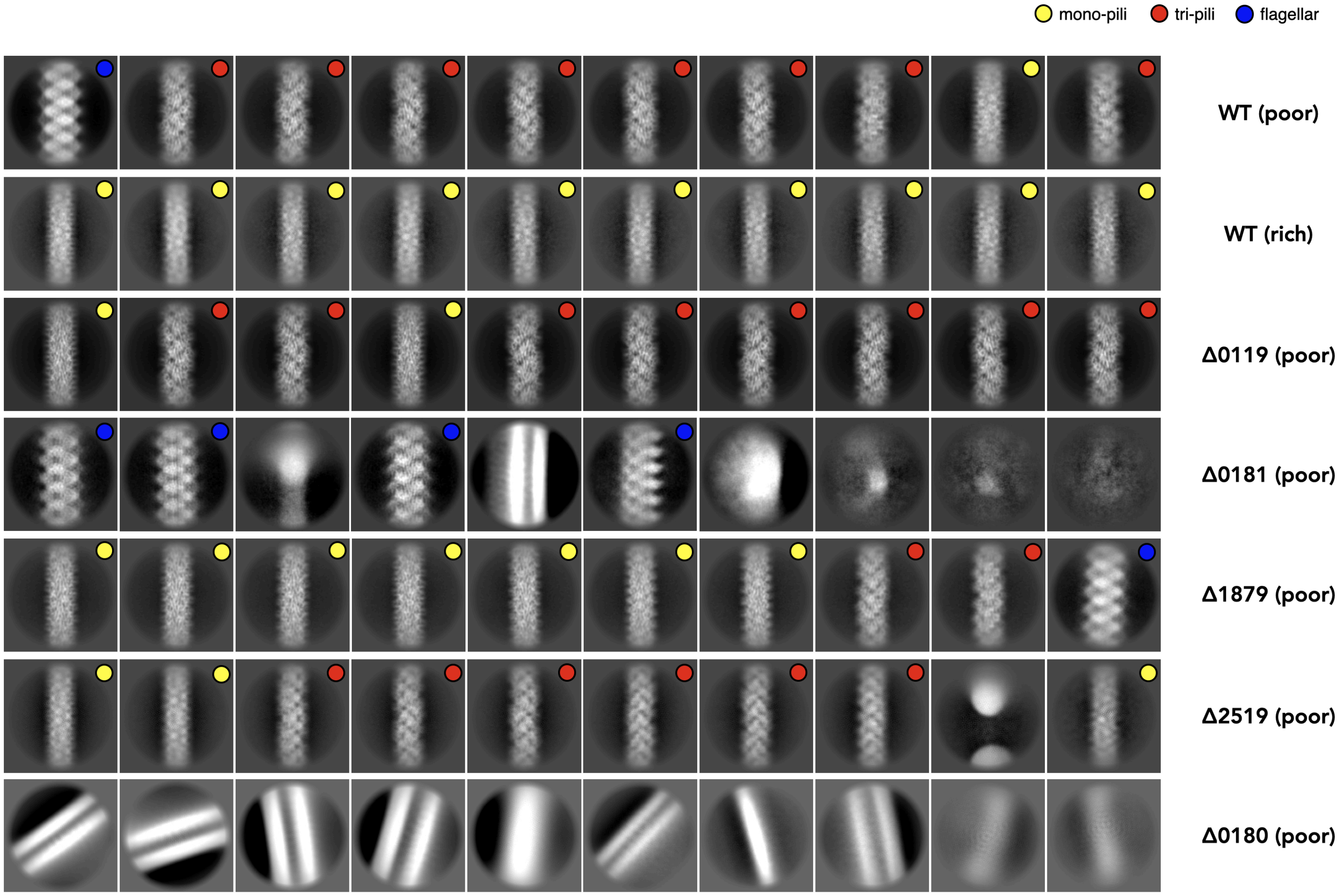


**Supplementary Figure 4. Pili production under different medium and ATPase knockout strains**
The top ten 2D classes for datasets collected in Fig 6c. The filament species identified based on the 2D averages, and their corresponding power spectrum are labeled on the upper right.

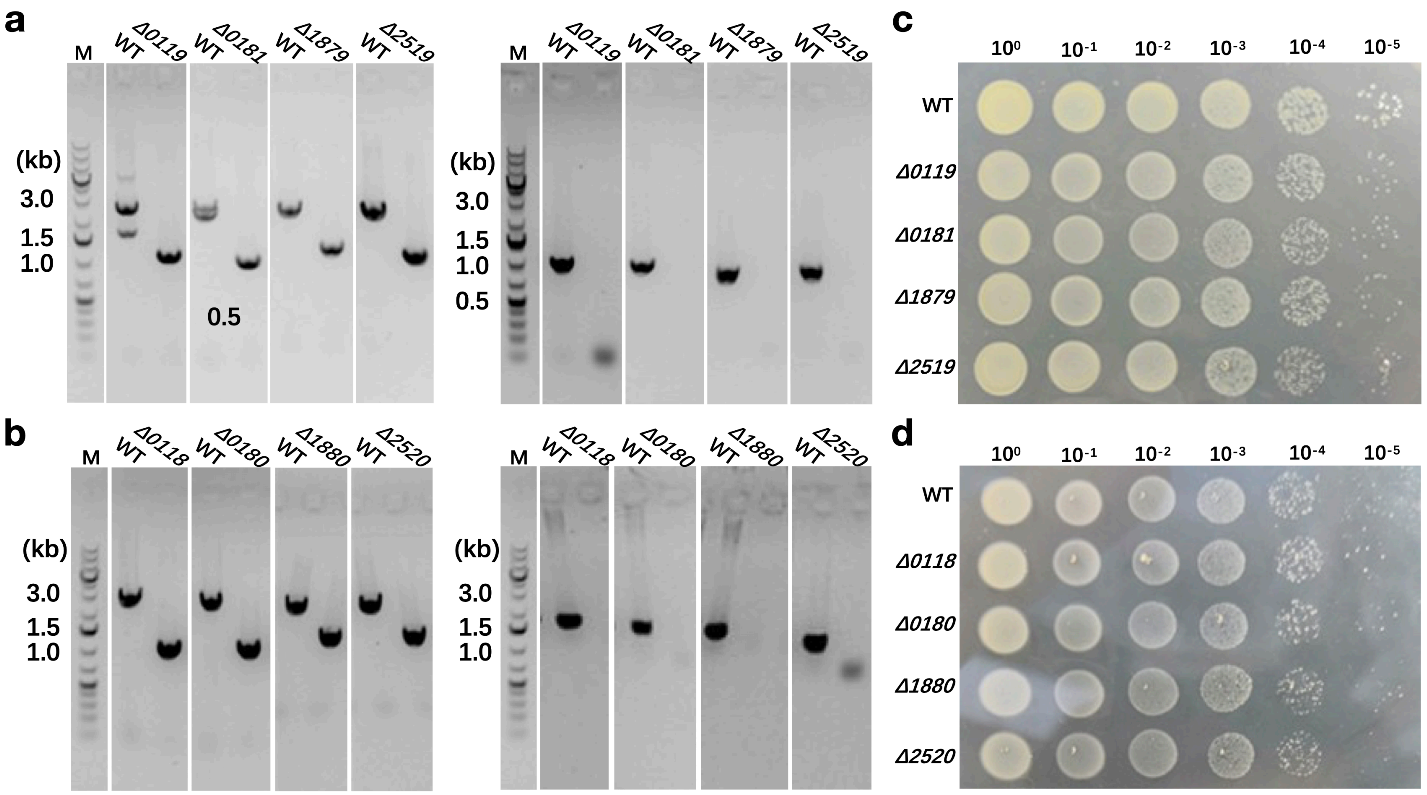


**Supplementary Figure 5. Construction of the ATPase and TadC knockout strains**

PCR verification of the ATPase (**a**) and TadC (**b**) knockout strains using the Flanking (Left) and gene specific (right) primers. Spot test to show the growth of the ATPase (**c**) and TadC (**d**) knockout strains. Knockout of the ATPase or TadC genes alone exhibited no obvious growth difference comparing with the WT strain on solid medium

### Supplemental Table 1. Plasmids used in this study

| **Plasmid** | **Description** | **Reference** |
|---|---|---|
| *pGE* | *Genome-editing plasmid* | Li et al[1] |
| pGE-*sire_0119* | *sire_0119* (ATPase gene) knockout | This study |
| pGE-*sire_0181* | *sire_0181* (ATPase gene) knockout | This study |
| pGE-*sire_1879* | *sire_0181* (ATPase gene) knockout | This study |
| pGE-*sire_2519* | *sire_2519* (ATPase gene) knockout | This study |
| pGE-*sire_0118* | *sire_0118* (TadC gene) knockout | This study |
| pGE-*sire_0180* | *sire_0180* (TadC gene) knockout | This study |
| pGE-*sire_1880* | *sire_1880* (TadC gene) knockout | This study |
| pGE-*sire_2520* | *sire_2520* (TadC gene) knockout | This study |

### Supplemental Table 2. Strains used in this study

| Strain | Phenotype | Reference |
|---|---|---|
| *S. islandicus REY15A* | *Wild type* | Guo et al[2] |
| Sis E2333 | REY15A *ΔpyrEFΔlacS* | Deng et al[3] |
| Sis/pGE | Control for knockout | Liu et al[4] |
| *Δsire_0119* | *sire_0119* knockout | This study |
| *Δsire_0181* | *sire_0181* knockout | This study |
| *Δsire_1879* | *sire_0181* knockout | This study |
| *Δsire_2519* | *sire_2519* knockout | This study |
| *Δsire_0118* | *sire_0118* knockout | This study |
| *Δsire_0180* | *sire_0180* knockout | This study |
| *Δsire_1880* | *sire_1880* knockout | This study |
| *Δsire_2520* | *sire_2520* knockout | This study |

### Supplemental Table 3. Spacers selected and used in this study

| Name | Sequence (5'-3') | Source |
|---|---|---|
| 0119-S-F | AAGCATACGAACAAGTTCATACCCTCGTCTAACATCATCCATA | This study |
| 0119-S-R | AGCTATGGATGATGTTAGACGAGGGTATGAACTTGTTCGTATG | This study |
| 0181-S-F | AAGAATGGCGATAGGATTGCCGCAACATTCAGACGTGAAGTAT | This study |
| 0181-S-R | AGCATACTTCACGTCTGAATGTTGCGGCAATCCTATCGCCATT | This study |
| 1879-S-F | AAGATAAGTCCTATATTATTGATTAAAAACGTATCGATAAGTT | This study |
| 1879-S-R | AGCAACTTATCGATACGTTTTTAATCAATAATATAGGACTTAT | This study |
| 2519-S-F | AAGGAAAAGCCTCTAACGATTATTGACTTGGTTTACAAGTATG | This study |
| 2519-S-R | AGCCATACTTGTAAACCAAGTCAATAATCGTTAGAGGCTTTTC | This study |
| 0118-S-F | AAGTCAGAGACTCTTTTAAGTTGCTTACTATCGCATAATTTCT | This study |
| 0118-S-R | AGCAGAAATTATGCGATAGTAAGCAACTTAAAAGAGTCTCTGA | This study |
| 0180-S-F | AAGTCTCTCTTTGGTTCAGTCCCATTTCCTATAATGGCTATTT | This study |
| 0180-S-R | AGCAAATAGCCATTATAGGAAATGGGACTGAACCAAAGAGAGA | This study |
| 1880-S-F | AAGATGATTTTACTGGCCTTTCAGTATATTTCGTCAACAATAA | This study |
| 1880-S-R | AGCTTATTGTTGACGAAATATACTGAAAGGCCAGTAAAATCAT | This study |
| 2520-S-F | AAGGTTTCGTTGGAGGTTTTACTCAGAAAGAACCTCATTCCTA | This study |
| 2520-S-R | AGCTAGGAATGAGGTTCTTTCTGAGTAAAACCTCCAACGAAAC | This study |

### Supplemental Table 4. Oligonucleotides used in this study

| Name | Sequence (5'-3') | Source |
|---|---|---|
| **Oligonucleotides used to amplify the L- and R-arm of the corresponding genes of interest** | | |
| 0119-L-arm-F-Sph I | AAGTACAATTGTGCT**GCATGC**CTATTAACCTAGCAGAATAA | This study |
| 0119-L-arm-R | TCCCTATCGAGGGCTTAACCTTCTTTATGACGAATTAAAC | This study |
| 0119-R-arm-F | GTTTAATTCGTCATAAAGAAGGTTAAGCCCTCGATAGGGA | This study |
| 0119-R-arm-R-Xho I | TAACATATTGGATG**CTCGAG**AAAAAGATGAAAGATGTAAA | This study |
| 0181-L-arm-F-Sph I | AAGTACAATTGTGCT**GCATGC**AGTAGAGTTTCCTTAATTTC | This study |
| 0181-L-arm-R | ATTGTTAATCTTGGTTTCAATATATATCACTTCTCCATCT | This study |
| 0181-R-arm-F | AGATGGAGAAGTGATATATATTGAAACCAAGATTAACAAT | This study |
| 0181-R-arm-R-Xho I | TAACATATTGGATG**CTCGAG**TACCCTAACTGATTTATATA | This study |
| 1879-L-arm-F-Sph I | AAGTACAATTGTGCT**GCATGC**GAGATGGGTATGATTTGGTC | This study |
| 1879-L-arm-R | ATTTCTGAAATTGGTATTTCTTCGCTTACTTTATAATAGC | This study |
| 1879-R-arm-F | GCTATTATAAAGTAAGCGAAGAAATACCAATTTCAGAAAT | This study |
| 1879-R-arm-R-Xho I | TAACATATTGGATG**CTCGAG**TAGAATTCCTTATTTCCTCG | This study |
| 2519-L-arm-F-Sph I | AAGTACAATTGTGCT**GCATGC**GTTGCAATAGCTGCGGTCAG | This study |
| 2519-L-arm-R | TCCTTACTCCAGCCTAATAACTCTAAGAGATATTATTCCC | This study |
| 2519-R-arm-F | GGGAATAATATCTCTTAGAGTTATTAGGCTGGAGTAAGGA | This study |
| 2519-R-arm-R-Xho I | TAACATATTGGATG**CTCGAG**CTATGAAACCTAATCTACTG | This study |
| 0118-L-arm-F-Sph I | AAGTACAATTGTGCT**GCATGC**GATGGTCTCAATTTTAGAGA | This study |
| 0118-L-arm-R | GATATGCCAAGAATATTTAGATATATTTACCTAAAATTAC | This study |
| 0118-R-arm-F | GTAATTTTAGGTAAATATATCTAAATATTCTTGGCATATC | This study |
| 0118-R-arm-R-Xho I | TAACATATTGGATG**CTCGAG**TGTAGCATTCCAAGCAATGC | This study |
| 0180-L-arm-F-Sph I | AAGTACAATTGTGCT**GCATGC**AAAAACCGTAAGGGATTATA | This study |
| 0180-L-arm-R | AGAGTGCCGAAATAGAGATTCCTATTGTTACTTTCTCAGA | This study |
| 0180-R-arm-F | TCTGAGAAAGTAACAATAGGAATCTCTATTTCGGCACTCT | This study |
| 0180-R-arm-R-Xho I | TAACATATTGGATG**CTCGAG**TCATTTACTGGTGGTTCCAA | This study |
| 1880-L-arm-F-Sph I | AAGTACAATTGTGCT**GCATGC**GACTATCATTAAGATATAGG | This study |
| 1880-L-arm-R | ATAATCAAATCTAAGTTACCTCAAATAGCCTTTGTCTCAA | This study |
| 1880-R-arm-F | TTGAGACAAAGGCTATTTGAGGTAACTTAGATTTGATTAT | This study |
| 1880-R-arm-R-Xho I | TAACATATTGGATG**CTCGAG**TACAATATTCTTAGTCCCAT | This study |
| 2520-L-arm-F-Sph I | AAGTACAATTGTGCT**GCATGC**GGGAAGTAAGAGGGAAAGAA | This study |
| 2520-L-arm-R | ATATGAACATTTCAGCTAGGCTTCTTTCAGTCACCCCTAT | This study |
| 2520-R-arm-F | ATAGGGGTGACTGAAAGAAGCCTAGCTGAAATGTTCATAT | This study |
| 2520-R-arm-R-Xho I | TAACATATTGGATG**CTCGAG**TCTGTCCGTTAAATGTGGTT | This study |
| **Oligonucleotides used to verify the knockout strains** | | |

| | | |
|---|---|---|
| 0119-Flanking-F | GGTAAGGGAGCTATTTAATGTAGC | This study |
| 0119-Flanking-R | CTACAGAACAAACTAGCAAAGACTA | This study |
| 0119-int-F | CTGCAGGATCATATGTAAATGCTGG | This study |
| 0119-int-R | CTAAGCAAGATAAGAATATCCGTACC | This study |
| 0181-Flanking-F | GATAGCCTCAGAGAGAATATTGGA | This study |
| 0181-Flanking-R | GATATTGTAGTTTATACCTTAGCCC | This study |
| 0181-int-F | GCCGAATCCTCACATCTTTATTACC | This study |
| 0181-int-R | CTCTTGTCCTCTTATCTCACCTAC | This study |
| 1879-Flanking-F | ACGAGATTTTCGAATGGAAGGAGTTA | This study |
| 1879-Flanking-R | GTAACATAGCCTCTCTTCTTAAACC | This study |
| 1879-int-F | GAGCTAACTGTTCCCTTAGCAGAT | This study |
| 1879-int-R | CACTGGCTTGAATCTTAGTTTACC | This study |
| 2519-Flanking-F | GGCTTAGAAAACGCTTTAGTTACTG | This study |
| 2519-Flanking-R | CTCTTAGTACGCTCGCTTATCCA | This study |
| 2519-int-F | CGAACCACAGCTTGATCAGAGAGA | This study |
| 2519-int-R | CATGACCACTGGCAACAGCTTGA | This study |
| 0118-flanking-F | GTCATGGCCCAGTTTACACTA | This study |
| 0118-flanking-R | GGAGAAATTAGAGATAGAGAAGG | This study |
| 0118-int-F | GTACTTTGCTTAACATGCTCAC | This study |
| 0118-int-R | GGAACTACTCTAATAACACAGC | This study |
| 0180-flanking-F | GGAATCTCTGGAATTAGGATAG | This study |
| 0180-flanking-R | GAAGAGGTAGAGAGAGAATTAC | This study |
| 0180-int-F | CACAGCAATGCGCTACGCTAG | This study |
| 0180-int-R | GTATTTGCTGTACACCACCATTC | This study |
| 1880-flanking-F | GCTAGACAATCAATCTCTTCCC | This study |
| 1880-flanking-R | CACGTTCTGAACAACACTATACG | This study |
| 1880-int-F | GTGGCGAGTCAATAGGAATTTC | This study |
| 1880-int-R | CATCAGCGACCGATATCTTAC | This study |
| 2520-flanking-F | CAGACTCTCCTTAAGATATAGAC | This study |
| 2520-flanking-R | GGTCCAATTACTATTGCATCTCC | This study |
| 2520-int-F | GTAAGAGACAGCATCCAAGATG | This study |
| 2520-int-R | CATATATCTAGGTCTATTGCTAGC | This study |
| **Oligonucleotides used to determine the transcriptional level of the corresponding genes by RT-qPCR** | | |
| 0119-qPCR-F | GCTACATTACCTTCTCTATCTC | This study |
| 0119-qPCR-R | CAGAAGTGACTAGAGAAACAG | This study |

| | | |
|---|---|---|
| 0181-qPCR-F | GTTCTTCAGTAGTGATACGAAG | This study |
| 0181-qPCR-R | GTAACTCCTATTGACATGATACT | This study |
| 1879-qPCR-F | GTCATAGGTTCTACCGGATC | This study |
| 1879-qPCR-R | CCTTATCCAGTTATCATGAACTA | This study |
| 2519-qPCR-F | GAAAGACCACTGCATTAAACTC | This study |
| 2519-qPCR-R | CATAGTTTGAAGGTCTAGTATAG | This study |
| 0118-qPCR-F | CTATTAGGAGTTGGTCTTAGG | This study |
| 0118-qPCR-R | CTTAGTACTGCCATCAGAGAC | This study |
| 0180-qPCR-F | GAAACTTTAGCTTCTACTGCTG | This study |
| 0180-qPCR-R | GCCATTATAGGAAATGGGACTG | This study |
| 1880-qPCR-F | GTCAATAGGAATTTCCATGCC | This study |
| 1880-qPCR-R | CCAGTAAAATCATTGGAGTGAC | This study |
| 2520-qPCR-F | CTCCACTTGACGCTTTAGAG | This study |
| 2520-qPCR-R | CAACGAAACTGGAGCCAATTAAG | This study |
| TBP-F | GTGGCAACAGTTACGTTAGAG | Liu et al[5] |
| TBP-R | CCTTGGGCTGTTCTAATCTG | Liu et al[5] |
